# Efficient Flexible Backbone Protein-Protein Docking for Challenging Targets

**DOI:** 10.1101/223511

**Authors:** Nicholas A. Marze, Shourya S. Roy Burman, William Sheffler, Jeffrey J. Gray

**Author notes:** These authors have equal contribution (Nicholas A. Marze, Shourya S. Roy Burman). The names are arranged alphabetically. Author for correspondence: Jeffrey J. Gray. Classification: BIOLOGICAL SCIENCES - Biophysics and Computational Biology.

## Abstract

Computational prediction of protein-protein complex structures facilitates a fundamental understanding of biological mechanisms and enables therapeutics design. Binding-induced conformational changes challenge all current computational docking algorithms by exponentially increasing the conformational space to be explored. To restrict this search to relevant space, some computational docking algorithms exploit the inherent flexibility of the protein monomers to simulate conformational selection from pre-generated ensembles. As the ensemble size expands with increased protein flexibility, these methods struggle with efficiency and high false positive rates. Here, we develop and benchmark a method that efficiently samples large conformational ensembles of flexible proteins and docks them using a novel, six-dimensional, coarse-grained score function. A strong discriminative ability allows an eight-fold higher enrichment of near-native candidate structures in the coarse-grained phase compared to a previous method. Further, the method adapts to the diversity of backbone conformations in the ensemble by modulating sampling rates. It samples 100 conformations each of the ligand and the receptor backbone while increasing computational time by only 20–80%. In a benchmark set of 88 proteins of varying degrees of flexibility, the expected success rate for blind predictions after resampling is 77% for rigid complexes, 49% for moderately flexible complexes, and 31% for highly flexible complexes. These success rates on flexible complexes are a substantial step forward from all existing methods. Additionally, for highly flexible proteins, we demonstrate that when a suitable conformer generation method exists, RosettaDock 4.0 can dock the complex successfully.

**Significance:** Predicting binding-induced conformational plasticity in protein backbones remains a principal challenge in computational protein–protein docking. To date, there are no methods that can reliably dock proteins that undergo more than 1 Å root-mean-squared-deviation of the backbones of the interface residues upon binding. Here, we present a method that samples backbone motions and scores conformations rapidly, obtaining–for the first time–successful docking of nearly 50% of flexible target complexes with backbone conformational change up to 2.2 Å RMSD. This method will be applicable to a broader range of protein docking problems, which in turn will help us understand biomolecular assembly and protein function.

## Introduction

Proteins bind each other in a highly specific and regulated manner. Often, a change in conformation from the unbound to the bound state forms the basis of the protein’s specificity and function in its interaction (1–4). Since the beginning of the field (5), conformational changes in proteins induced by binding have confounded protein–protein docking algorithms by greatly increasing the degrees of freedom to be sampled. While rotamer libraries have alleviated the sampling challenges for surface side chains (6), backbone flexibility remains the principal challenge in protein–protein docking. Previous studies have found limited success by varying the backbone along a restricted set of coordinates (7–9) or interface residues (10, 11) or by docking a small number of backbone conformations of the two partners (12–15). The most recent rounds of the blind docking challenge, Critical Assessment of PRediction of Interactions (CAPRI), demonstrated that protein flexibility is still a community-wide weakness, with flexible target complexes eliciting no successful predictions from any method (16, 17).

Flexible-backbone docking, as well as other key remaining protein–protein docking challenges such as global docking and docking of large multi-domain complexes, demands more algorithmic complexity to explore a larger conformational search space than rigid-body docking of small proteins (18). Coarse-graining is commonly used to model longer time-scales and larger systems in a rapid, yet meaningful manner (19, 20). Score functions designed to navigate this reduced space smoothen the energy landscape to avoid getting stuck in local minima. While allowing orders-of-magnitude more conformational sampling, coarse-grained models are limited by their accuracy and typically require refinement in higher resolution stages.

The consensus on the kinetic mechanism of most of these conformational changes is that the protein monomers exist in an equilibrium of multiple conformations from which the preferred conformations are selected during an initial encounter with the binding partner, and subsequently, localized structural rearrangements stimulated by the partner tightens the binding (21, 22). The former mechanism is called conformational selection, and it lends itself to coarse-graining as the discrete conformations can be individually sampled. However, large conformational ensembles of flexible proteins multiply the computational demand and increase the false positive rates. Previous studies have used experimental data to create a minimal ensemble that captures the observed flexibility (23), but these data are seldom available. Thus, it is desirable to have a coarse-grained method that efficiently samples a sizeable ensemble while distinguishing spurious interfaces from the native interface. Smaller changes caused by induced fit are less suitable to be modeled at this resolution, but are more amenable to full-atom modeling.

RosettaDock has historically been among the top-performing methods for computational protein–protein docking (24–28). Combining coarse-grained conformational selection with full-atom induced fit, RosettaDock 3.2 achieved successful docking predictions on a majority of rigid complexes (58%) in the Docking Benchmark 3.0 set (29). On the more flexible targets, however, RosettaDock (like other methods) performed poorly, only achieving a successful docking prediction on 29% of the moderately flexible complexes and 14% of the highly flexible complexes. The performance in CAPRI rounds since the last major version release mimicked the benchmark performance (30). For flexible docking, the current protocol relies on sampling a pre-generated ensemble of monomer backbone conformations (13), but increasing the ensemble size beyond 20 conformers is computationally infeasible. Additionally, the “centroid” score function used to discriminate near-native conformations from incorrect ones is not sufficiently accurate in the coarse-grained phase, where the search is the broadest (31).

In this study, we pursued two avenues to address these computational limitations. First, to improve sampling efficiency, we developed a fast and scalable backbone sampling algorithm, Adaptive Conformer Selection (ACS), that modulates the frequency of conformer selection for each partner depending on the size and diversity of the ensemble. Second, to improve scoring efficiency, we developed a fast and accurate scoring method, Motif Dock Score (MDS), based on the residue-pair transform (RPX) score, which was recently developed and used to design hydrophobic symmetric protein interfaces (32). RPX score evaluates and mutates residue pairs using the 6D transformation needed to superimpose the residues’ N–C_α_–C backbone atoms onto each other. In a single lookup, RPX score queries this transformation against a pre-tabulated database of aliphatic amino acid pairs and their corresponding geometries and full-atom Rosetta scores. The pair score and sequence of the best amino acid pair from the database are then assigned to the queried residue pair. We derived and optimized MDS from the RPX basis in the context of the RosettaDock protocol, expanding it to all twenty amino acids and selecting for enrichment of near-native candidate structures.

We tested RosettaDock 4.0, which contains both ACS and MDS enhancements, on a subset of Docking Benchmark 5.0 (4) to evaluate the relative performance versus RosettaDock 3.2, and other commonly used docking protocols. The performance in both the full benchmark set and the three flexibility-based subsets (rigid, medium-flexible, and highly flexible) showed significant improvements, most notably among previously intractable flexible-backbone complexes.

## Results

### Adaptive Conformer Selection

RosettaDock is a Monte Carlo-plus-minimization algorithm (33) consisting of a low-resolution stage, which simulates conformer selection during the formation of the encounter complex, followed by a high-resolution stage, which simulates induced fit in the bound complex (13, 34). To produce a variety of starting states for the different trajectories, the ligand (the smaller protein) is first randomly rotated and translated about the receptor (the larger protein). In the low-resolution stage, side chains are replaced by coarse-grained “pseudoatoms”, allowing the ligand to efficiently sample the interface by rigid-body movements in a smoothened energy landscape. These rigidbody moves are coupled with backbone conformation swaps where the current backbone conformations of the ligand and the receptor are swapped with different ones from a pre-generated ensemble of conformations. In the high-resolution stage, the side chains are reintroduced to the putative encounter complex and those at the interface are packed for tight binding. There is minimal rigid-body motion in this second stage.

The previous version of RosettaDock, version 3.2, was optimized to handle smaller ensembles and hence had a fixed number conformation swaps. This choice led to reduced sampling of near-bound conformations as the ensembles grew larger. In RosettaDock 4.0, we alleviate this problem by modulating the number of conformer swaps depending on the swap acceptance rate of the previous cycle. If the acceptance rate of the conformer swaps is under 30%, the ensemble is presumed to be large and diverse, and hence the probability of the conformer swap is increased by 25%; conversely, if the acceptance rate is 30% or more, the probability is reduced by 25%. This adjustment helps prevent unnecessary backbone sampling for small ensembles and those with similar backbones while increasing backbone sampling for diverse ensembles by up to 477% over the course of 8 cycles. We call this backbone variation method Adaptive Conformer Selection (ACS). Fig. 1A shows the variation in conformer sampling frequencies for an example case of the ClpA chaperone:Clp protease adapter complex (PDB: 1R6Q), where the unbound to bound deviation of the C_α_ atoms at the interface is 1.4 Å for the chaperone and 2.0 Å for the protease. In this case, the protocol adapts to enable more trials of the protease backbone conformer swaps, and to a lesser effect the chaperone too.

**Fig. 1.**
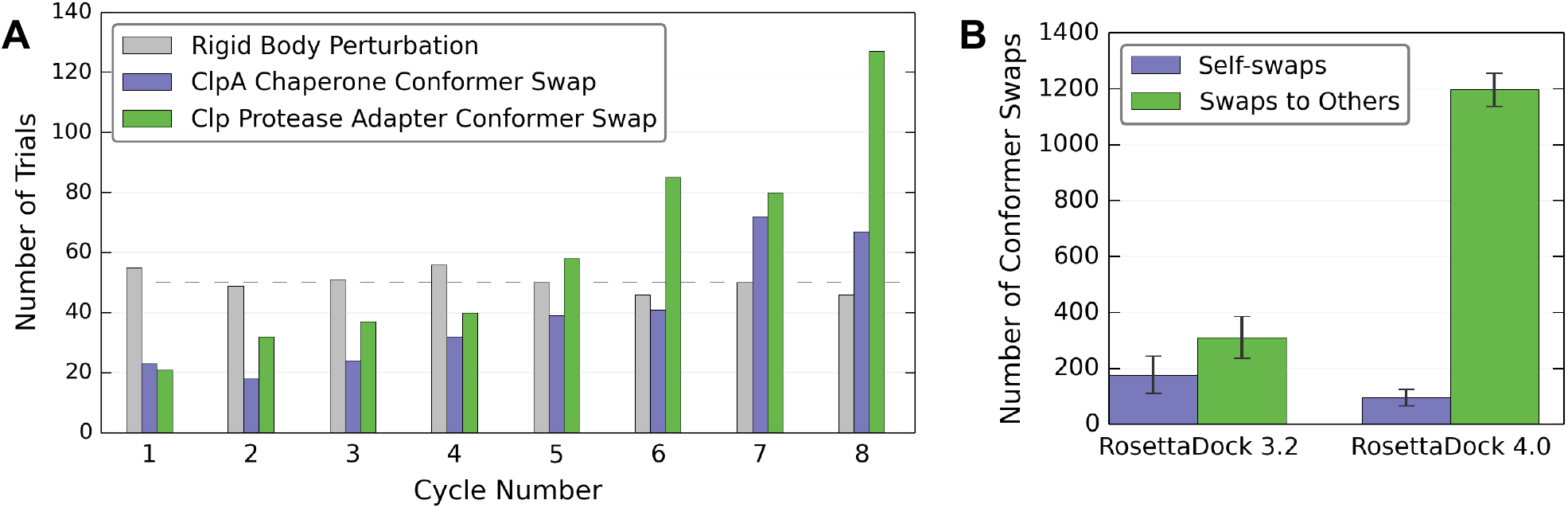
Amount of backbone sampling in RosettaDock 4.0. (A) Modulation of backbone conformer swap trials in Rosetta 4.0 for each of the first 8 cycles of Monte Carlo moves in the low-resolution search stage. The dashed line indicates the number of trials for each of the different moves in RosettaDock 3.2. Adaptive conformer selection in RosettaDock 4.0 ensures increased backbone swapping frequency for Clp protease adapter over ClpA chaperone, which is less flexible at the interface. (B) Comparison of the number of self-swaps versus swaps to other conformations in RosettaDock 3.2 versus Rosetta 4.0 for the highly flexible CCS metallochaperone: superoxide dismutase complex. RosettaDock 4.0 has increased backbone sampling both in the number and fraction of other conformations sampled.

**Fig. 2.**
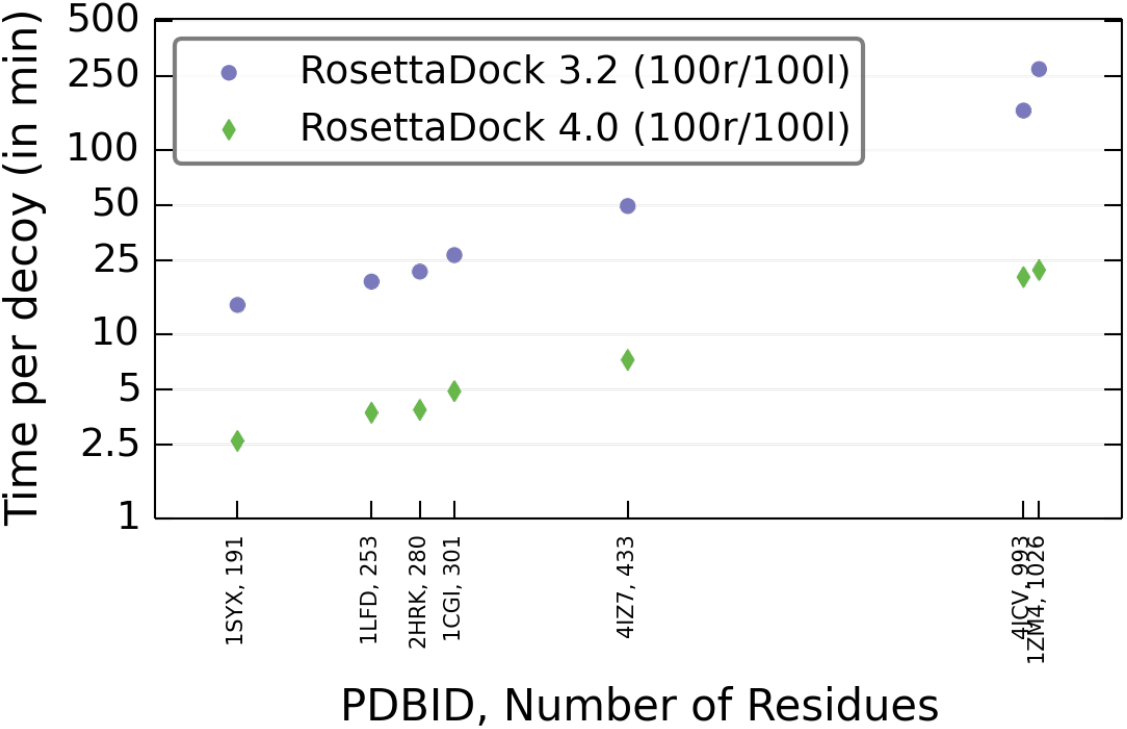
Time comparison of the docking protocols for large ensembles. Average time per decoy for RosettaDock 3.2 (blue) and 4.0 (green) with ensembles having 100 receptor and 100 ligand conformations for complexes ranging from 191 to 1026 total residues. Adaptive Conformer Sampling makes RosettaDock 4.0 up to 12 times faster for cases with large interfaces.

Previously, to determine which backbone was to be swapped in during conformer swapping, RosettaDock calculated the partition function of the entire ensemble of backbones superimposed along the protein–protein interface. The constraints of the interface, steric and otherwise, penalized conformations with backbone variations near the interface, creating a high probability for the existing backbone to be reselected during the conformer swap. In the case of superoxide dismutase (PDB: 1JK9), 36% of the backbone swaps were self-swaps (Fig 1B). Moreover, if there are *n_1_* conformations of the receptor and n_2_ conformations of the ligand, the partition function calculation required *O*(*n_1_*•*n_2_*) time, which meant that it required 10^3^ times longer for ensembles with 100 conformations each than for ensembles with 1 receptor conformation and 10 ligand conformations (13). We replaced this expensive partition function calculation with random conformer swaps, speeding up the protocol by as much as 12-fold and reducing self-swapping to 8% (approximately the inverse of the size of the ensemble). Instead of simultaneously swapping backbones of both partners and making a rigid-body move, as was done in the earlier protocol, we randomized the order of the receptor backbone swap, the ligand backbone swap and the rigid-body move.

#### Efficiency of Conformer Sampling

ACS made RosettaDock 4.0 marginally faster than RosettaDock 3.2 for simulations with small ensembles of 1 receptor and 10 ligand conformations (Fig S1). The speed-up was pronounced when the ensembles of both partners have 100 conformations each. For protein complexes larger than 1000 total residues, for example, eEF2-ETA-bTAD complex (PDB: 1ZM4) with 204 residues in the ligand and 822 residues in the receptor, ACS was over 12 times faster than RosettaDock 3.2 (Fig 2). Thus, the ACS method scales up practically for larger ensembles.

### Optimization of Motif Dock Score

For the recognition of the native interface during the broad, low-resolution search, docking requires a score function with predictive accuracy close to that of the well-tested full-atom score function (35). In earlier versions of RosettaDock, the low-resolution “centroid” score function relied on a single distance between potential interacting residues to score inter-chain contacts. This one-dimensional information was insufficient to represent the relative orientation of the two residues and consequently, their interaction. A statistical potential derived by using two interresidue distances (C_α_–C_α_ and C_β_–C_β_) showed remarkable accuracy on Bcl-2 affinity predictions (36), suggesting that with more information on relative orientation, it could be possible to distinguish native interfaces without representing the side chain in full. With this idea in mind, we developed Motif Dock Score (MDS) based on the residue-pair transform (RPX) framework (32) for interface design. MDS calculates the 6-dimensional transform (3 rotations and 3 translations) needed to superimpose the backbone atoms of interacting residues, looks up the residue pair score from pre-generated tables, and sums scores over all such pairs. Each entry in these tables is the lowest full-atom score calculated for a pair of interface residues in the bin for the given relative backbone orientation.

MDS depends on a discrete space tabulation of all-atom energies; therefore, the bin size of the scoring grid affects its performance. For our first round of optimization, we examined the relative performance of five bin sizes, ranging from 8 Å to 0.5 Å for translations and from 36° to 12° for rotations. We built experimental scoring grids for each of the five bin sizes and evaluated their performance by rescoring decoys from a set of eleven test targets (set 1, see Methods). As shown in Fig. 3A, both the largest and smallest bins performed poorly. The largest bin size convolutes too many distinct geometries into a single bin, compromising its ability to discriminate near-native structures from non-native structures. Conversely, structures scored by the smallest bin size typically have fewer than 20 non-zero pairwise interactions as they are too sparsely populated to score most pair interactions. We selected the 2 Å/22.5° bin size as optimal. We also tested smoothing the scoring grid for optimization, but it had little effect on the scoring performance (Supplementary Method S1 and Figs. S3 and S4).

**Fig. 3.**
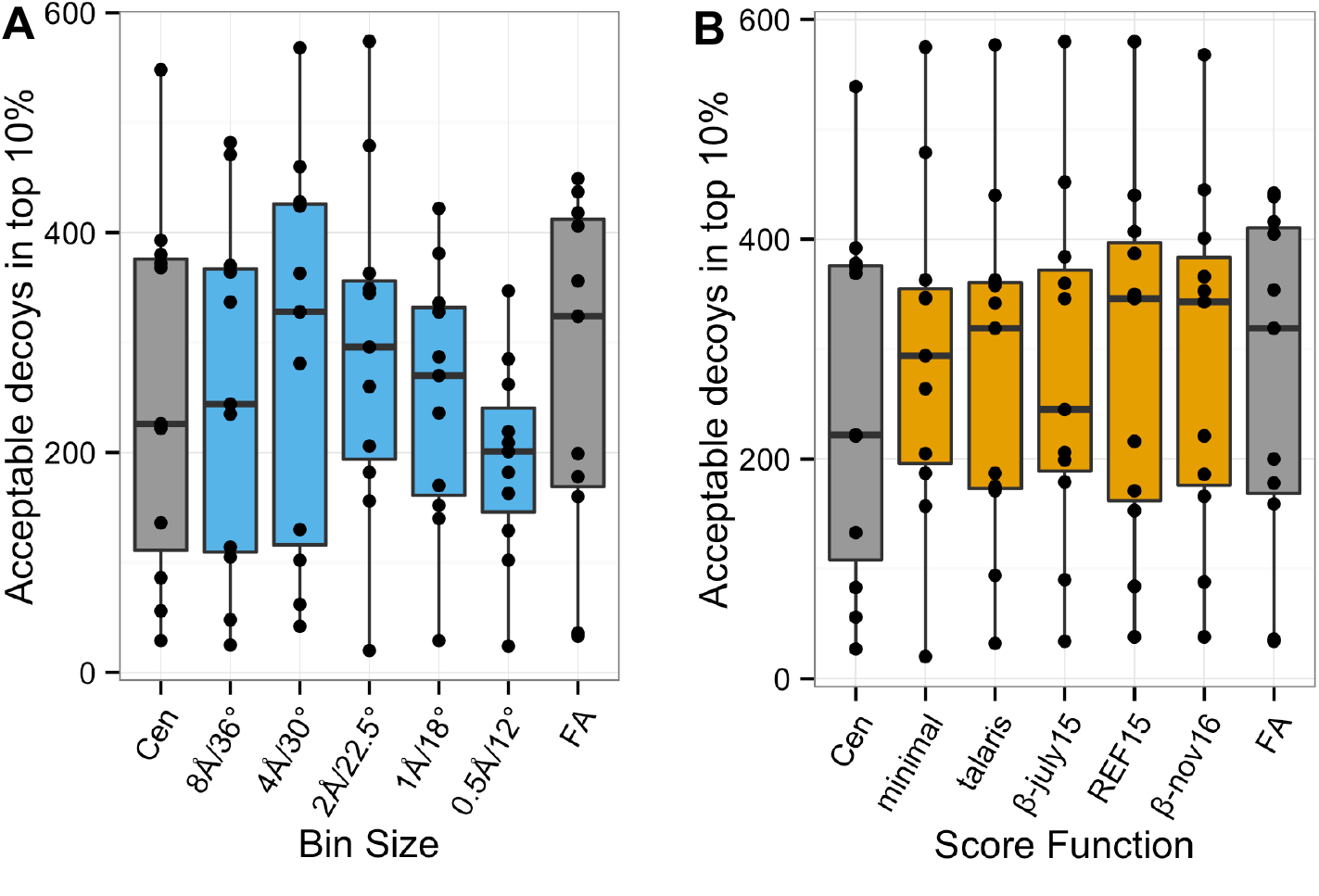
Optimization of bin size and underlying score function for motif dock score. Near-native structure enrichment for one representative MDS score table formulation for (A) each of the five examined bin sizes, and (B) four different motif-generating score functions. Each formulation trial is represented by a boxplot showing the enrichment of CAPRI-rating acceptable models within the 1,000 lowest-scoring models out of a set of 10,000 (top 10%) for each of 11 protein–protein complexes. The enrichment performance for RosettaDock’s original centroid scoring method (Cen) and the full-atom scoring method (FA) are shown in gray for comparison. (A) The intermediate bin sizes show the best near-native enrichment. (B) Four score functions are examined in comparison to the RPX default *minimal* score function: *talaris2014*, *beta_july15*, *REF15*, and *beta_nov16*. The score functions are examined absent their single-body score terms. The *talaris2014, REF15*, and *beta_nov16* score functions all outperform the minimal score function, with the latter two slightly outperforming even the full-atom score function in regards to near-native enrichment.

For our second round of optimization, we tested alternate underlying score functions to generate the residue pair motifs. Initially, we built motif tables from our PDB set using only the two-body terms (detailed formulation in Supplementary Method S1) of four score functions: *talaris2014*, the former Rosetta standard (37), and three iterations of the current Rosetta standard, *REF15* (35, 38), *betajulyl5* and *beta_nov16* (39). From these motif tables, we built score grids with the optimal bin size and rescored our docking test set. Among the four score functions, the *REF15* score function had the highest average near-native enrichment of all score grids tested (Fig. 3B). The *REF15* is also the most recent stable, released version of the score function. Finally, to prevent protein partners from embedding in each other, a van der Waals repulsive term (*interchain_vdw*) defined for the low-resolution mode was added to our final Motif Dock score function (see Fig. S5).

### Benchmarking Motif Dock Score

#### Evaluation Metrics

We evaluated the results of the docking benchmark runs using two types of metrics: a top-scoring near-native model count and near-native enrichment values. We define *N#* as the number of near-native decoys among a set number (#) of top-scoring decoys after the high-resolution stage, analogous to the *N5* metric used in previous studies (29). An expected value for *N#* metrics, 〈*N#*〉, can be calculated by a resampling strategy that helps quantify the stochasticity in decoy structure generation (see Methods). Enrichment values measure how the fraction of near native structures are enriched in a set of the top N% scoring decoys after the low-resolution stage. *E_N_%* is defined as:

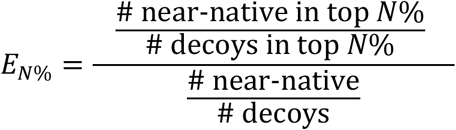

#### Benchmark Performance

To evaluate the accuracy of local docking using MDS, we compared its performance against a baseline method, RosettaDock 3.2’s centroid low-resolution docking mode, on a representative, nine-target benchmark set (set 2, see Methods). For each of the two algorithms, we generated 10,000 candidate structures per complex. As examples, Fig. 4 shows the Ras:RALGDS domain complex (PDBID: 1LFD) and BET3:TPC6 complex (PDBID: 2CFH) results. All candidate structures generated by the low-resolution phase of docking are plotted, comparing their low-resolution score to their RMSD from the experimental bound structure. For the baseline score function (Fig. 4A), the lowest-scoring models are nearly all incorrect with RMSD values from 7 to 22 Å, and few models under 5 Å are sampled at all. In contrast, with MDS (Fig. 4B), a clear "funnel" can be seen in the plot, with the lowest-scoring models having low RMSD values from the native structure. The top-scoring structures are near-native indicating a successful discrimination. Further, if MDS was used to filter the candidate structures so that only the top 1 or 10% of low-resolution candidates were sent to the computationally intensive refinement stage, near-native structures would be included in the set. In contrast, filtering with the centroid score would eliminate the best structures. Docking results of BET3:TPC6 complex (Fig. 4C and 4D) present a similar trend in that near-native models are lost when filtering on centroid score and can be retained by filtering on MDS. Table S2 presents docking metrics for each of the nine complexes in the test set. Significant improvements occur for all but the most flexible complexes.

**Fig. 4.**
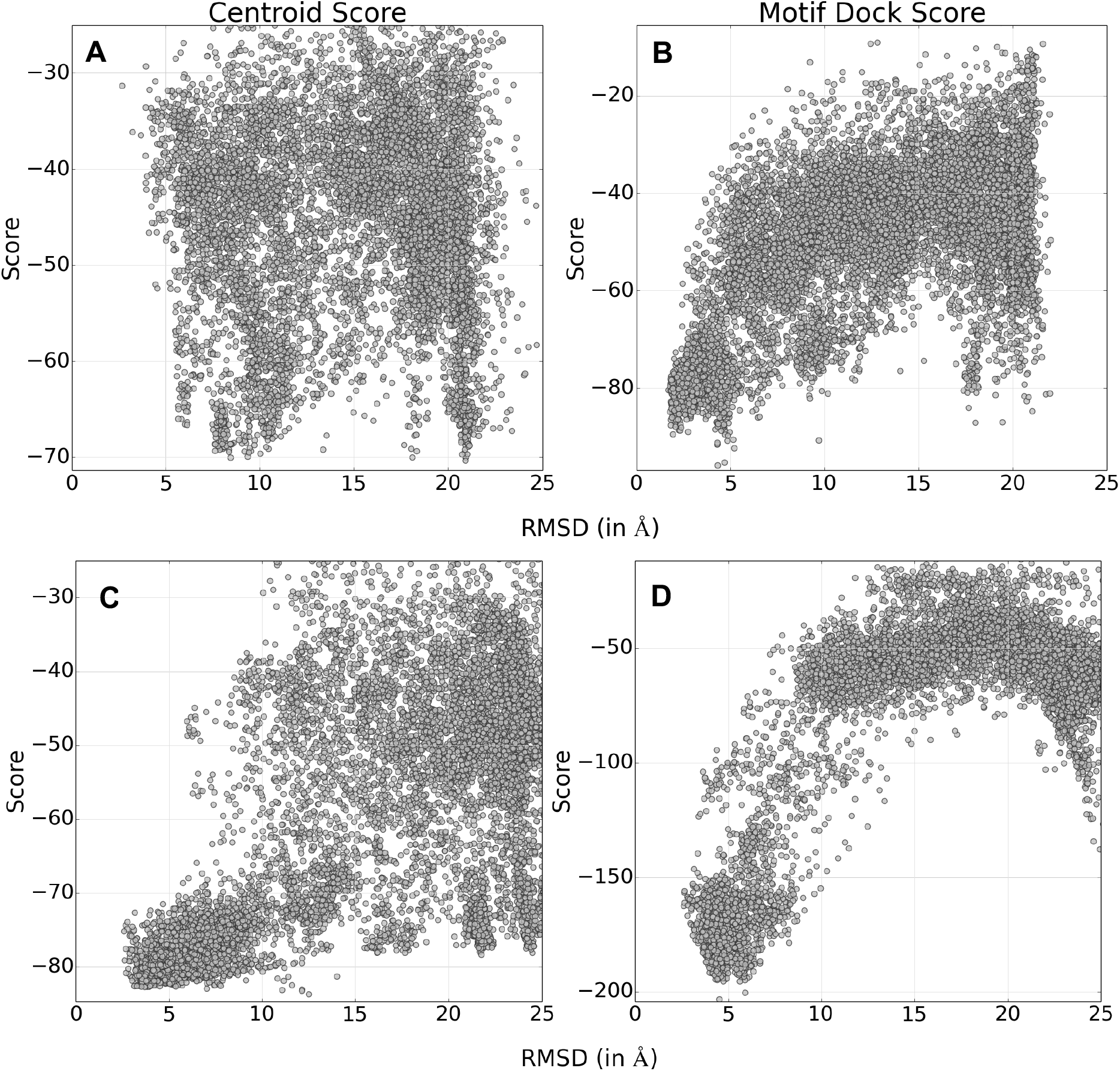
Low-resolution score vs. RMSD from native plots for two examples. *viz*. Ras:RALGDS domain complex (A and B) and BET3:TPC6 complex (C and D). (A and C) 10,000 models generated by RosettaDock 3.2 using the centroid score function, and (B and D) 10,000 models generated by RosettaDock 4.0 using motif dock score (MDS) function. (A) Centroid score does not generate many near-native candidate structures, and it cannot distinguish them from incorrect models. All metrics indicate failure: N5 = 0, N100 = 0, N1000 = 23. (B) MDS generates a large number of near-native candidate structures, and discriminates them from incorrect models. All metrics indicate success: N5 = 5, N100 = 95, N1000 = 750. (C) N5 = 1 indicates discrimination failure, but N100 = 86 and N1000 = 673 indicate that the broader set is enriched in near-native structures. (D) All metrics indicate success: N5 = 5, N100 = 98, N1000 = 813.

To test whether MDS was unduly biased by existing structures of homologous interfaces in creating the score function, we removed all homologs of the proteins in Docking Benchmark 5.0 identified in the Dockground (40) and the PIFACE (41) libraries before building the motif tables. Table S3 demonstrates that the performance of MDS with tables built after removal of the 8,126 homologs is similar to that with just the benchmark PDBs removed.

Encouraged by the promise that MDS showed in local docking simulations, we also tested MDS in global docking simulations. Unfortunately, 10,000 docking decoys were insufficient to produce thorough global sampling, with no more than a handful of sub-10 Å structures generated for any target (Fig. S7). The results show, however, that global docking with MDS recapitulates the energy landscape observed in local docking, and that four targets have no well populated, lower-scoring, false energy funnels, suggesting that a standard MDS global docking run would produce and enrich near-native structures in many targets.

### Advantage of Using Large and Varied Ensembles

In a blind prediction, where the location and extent of backbone motions is unknown, an ensemble generated using multiple conformation generation methods is more likely to contain a near-bound conformation than one generated from a single source. To delineate the gains made by using larger and more varied ensembles from the method improvements, we tested the two protocols, RosettaDock 3.2 and RosettaDock 4.0 with both small, similar ensembles and large, diverse ensembles. The small ensembles contained 1 receptor and 10 ligand conformations, all of which were made using Rosetta’s Relax protocol (42). The large ensembles contained a mixture of conformations generated using Relax, Backrub (43) and normal modes analysis (NMA) (44) for a total of 100 conformations of each docking partner.

The results of Ras:RALGDS domain complex (PDB: 1LFD) are depicted in Fig. 5, where a large loop motion (RMSD_Cα_ of 2.2 Å) helps the RALGDS domain interact with Ras. With RosettaDock 3.2 and the smaller ensembles (Fig. 5A), few models could be classified as medium-quality, but, more critically, models with false interfaces had lower scores than these models, rendering the simulation unsuccessful on the *N5* metric. Docking with RosettaDock 4.0 and the smaller ensembles (Fig 5B) showed an increase in enrichment of medium-quality models and a successful dock. However, the RALGDS domain in the best scoring acceptable structures was rotated to enable the loop interact with Ras (Fig. 5E). Using the larger ensembles, both RosettaDock 3.2 (Fig. 5C) and 4.0 (Fig. 5D) find deeper funnels due to the presence of conformations generated using the Backrub protocol where the contacting residues in the loop were within 0.2 Å of the bound structure. While RosettaDock 3.2 recognizes these near-bound conformations, they are sampled infrequently. This distribution qualifies as a success on the *N5* metric, but the near-native enrichment was low. Moreover, it took an average of 19.3 minutes to generate a structure as opposed to 3.7 minutes for RosettaDock 4.0. With the protocol improvements in RosettaDock 4.0 and a large ensemble to sample, the best scoring models recovered up to 64% of the native residue-residue contacts (Fig. 5F).

**Fig. 5.**
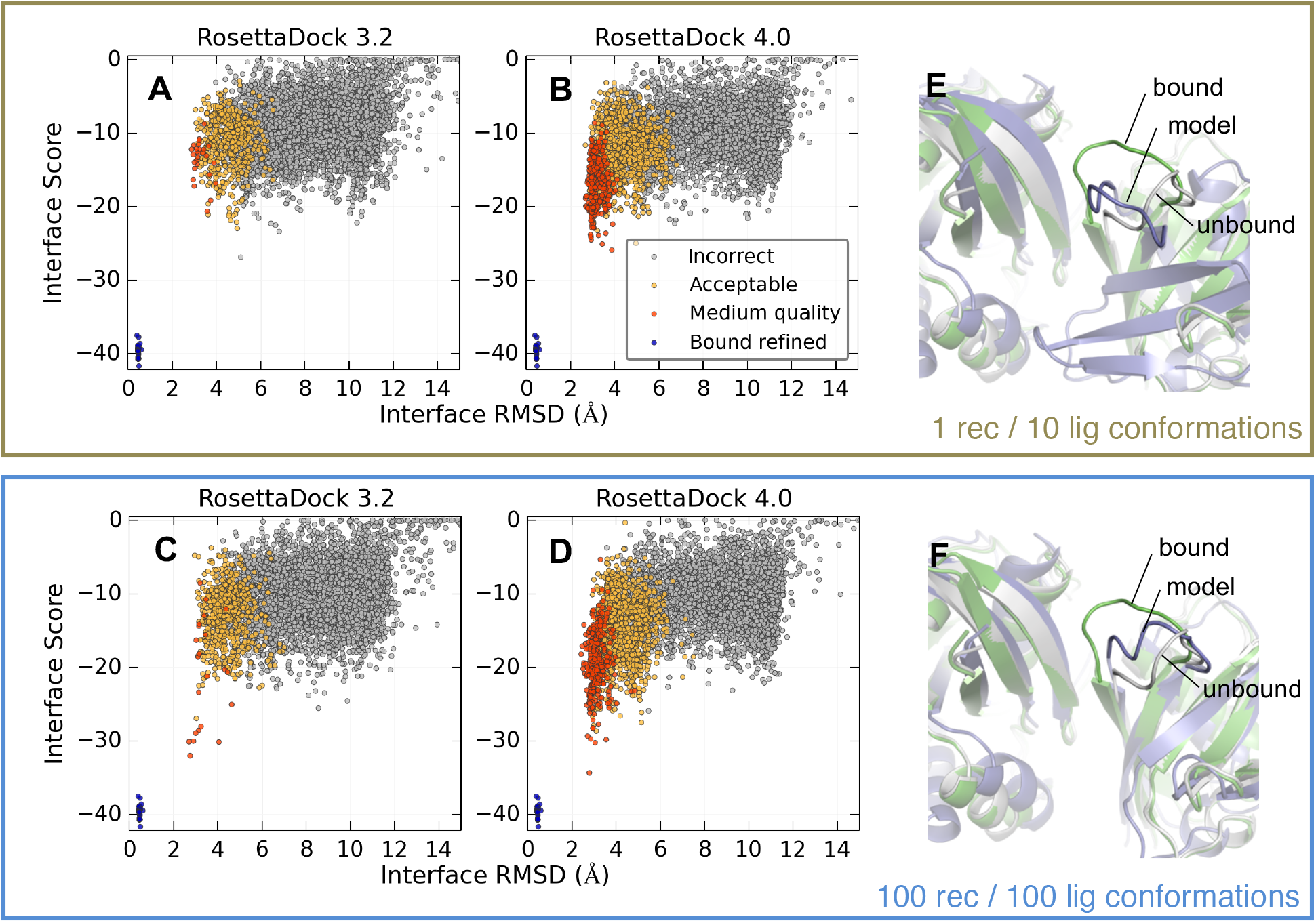
Comparison of docking protocols on Ras:RALGDS domain complex with different backbone ensembles. Interface score versus interface RMSD plots for docking simulation with (A) RosettaDock 3.2 and ensembles with 1 receptor and 10 ligand conformations generated bythe Relax protocol, (B) RosettaDock 4.0 and ensembles with 1 receptor and 10 ligand conformations generated by the Relax protocol, (C) RosettaDock 3.2 and ensembles with 100 conformations each of the receptor and the ligand generated by the Relax, Backrub and NMA protocols, and (D) RosettaDock 4.0 and ensembles with 100 conformations each of the receptor and the ligand generated by the Relax, Backrub and NMA protocols. Colored points indicate CAPRI-quality category for each decoy, and the blue points provide a reference energy of the refined, bound crystal structure. (B) and (D) are enriched in medium-quality docked models as compared to (A) and (C), respectively. (C) has a deeper funnel than (A) owing to the inclusion of conformations generated by Backrub, which produces loop motions that mimic the unbound-bound conformational change. (D) has both a deep funnel and enhanced sampling. (E) The best docked structure (in blue) for runs with the smaller ensemble has the RALGDS domain rotated to find the interface interactions. (F) The best docked structure (in blue) for runs with the larger ensemble has a better overall RMSD and f_na_t recovery. The superimposed unbound structures are in white and the bound structures are in green.

A similar phenomenon is observed in the case of glutamyl-tRNA synthetase:GU4 nucleic-binding protein 1 complex (PDB: 2HRK) where the interface undergoes a collective motion of 1.8 Å RMSD. A small, collective motion in the conformations generated by NMA prevents the backbone atoms of Asn-124 in glutamyl-tRNA synthetase from clashing with those of Arg-102 on GU4 nucleic-binding protein, allowing for a tighter interface (Fig. S8). These examples suggest that swift and adequate sampling of large ensembles generated from different sources better produces native-like interfaces.

### Evaluation of RosettaDock 4.0 on Benchmark Set

The ensemble generation methods used, *viz*. Rosetta Backrub, Rosetta Relax and NMA, have been shown to produce backbones that are between 1 and 4 Å RMSD from the unbound starting structure, with an average correlation of 0.4-0.5 to the experimentally determined displacements of the bound and unbound states (18). The extent of motion suggested that the ensembles generated using these methods could be used to dock moderately flexible proteins. Thus, we built a benchmark set heavily enriched with moderately flexible proteins to evaluate the RosettaDock 4.0 protocol.

We evaluated the accuracy of RosettaDock 4.0 for 43 complexes classified as medium-flexible, as well as for 32 classified as flexible and 13 classified as rigid, for a total of 88 targets (set 3, see Methods). For each target, we pre-generated 100 conformations for both the ligand and the receptor ensembles. The three conformer generation methods produce motions in different directions and locations, and hence we increased the variability of the full ensembles by using 40 conformations made using NMA, 30 made using Backrub and 30 made using Relax. We then generated 5,000 local docked models using the full RosettaDock 4.0 protocol for each target. We also ran control simulations using the RosettaDock 3.2 protocol, also generating 5,000 candidate structures per target. For a fair comparison to the previously published accuracy metrics, we generated conformer ensembles for the control runs containing only 1 receptor conformation and 10 ligand conformations.

The ability of the two protocols to sample and discriminate near-native structures was evaluated using the bootstrapped *N5* average, 〈*N5*〉, both after the low-resolution stage and for the final models after the high-resolution stage. To evaluate the enrichment in the low-resolution stage alone, which dictates how many trajectories need to be run, we used the 〈*E_1%_*〉 metric. As summarized in Table 1, RosettaDock 4.0 shows significant performance gains over RosettaDock 3.2, particularly in the low-resolution phase. RosettaDock 4.0’s near-native enrichment is improved markedly, with median 〈*E_1%_*〉 of 2.5, implying that its very low-scoring sets are significantly enriched with near-native structures from the bulk candidate set. RosettaDock 3.2’s median 〈*E_1%_*〉 value is 0.0, indicating that the very low-scoring set is devoid of near-native structures. Fig. 6 compares enrichments of RosettaDock 3.2 versus RosettaDock 4.0 for each target. The 〈*E_1%_*〉 performance (Fig. 6A) improves for 62 complexes in RosettaDock 4.0, most of which had zero enrichment previously. The performance is worse for 7 complexes, primarily due to favorable scoring of spurious interfaces. For the remaining 19 complexes, neither method was enriched in near-native decoys. RosettaDock 4.0 has an average low-resolution 〈*N5*〉 value of 1.3 across all targets, which implies that even after coarse-graining the side chains, more than one in the five top-scoring structures is near-native on average. This is approximately a ten-fold improvement over the corresponding average from RosettaDock 3.2. Our criterion for success discrimination is that the 〈*N5*〉 value should be 3 or higher. We see a seven-fold improvement in the number of expected low-resolution discrimination successes across the benchmark set (16.8 vs. 2.5 complexes). Pairwise target comparison (Fig. 6B) shows that only 2 success cases are lost from RosettaDock 3.2 to RosettaDock 4.0, while 13 additional successes are added.

**Fig. 6.**
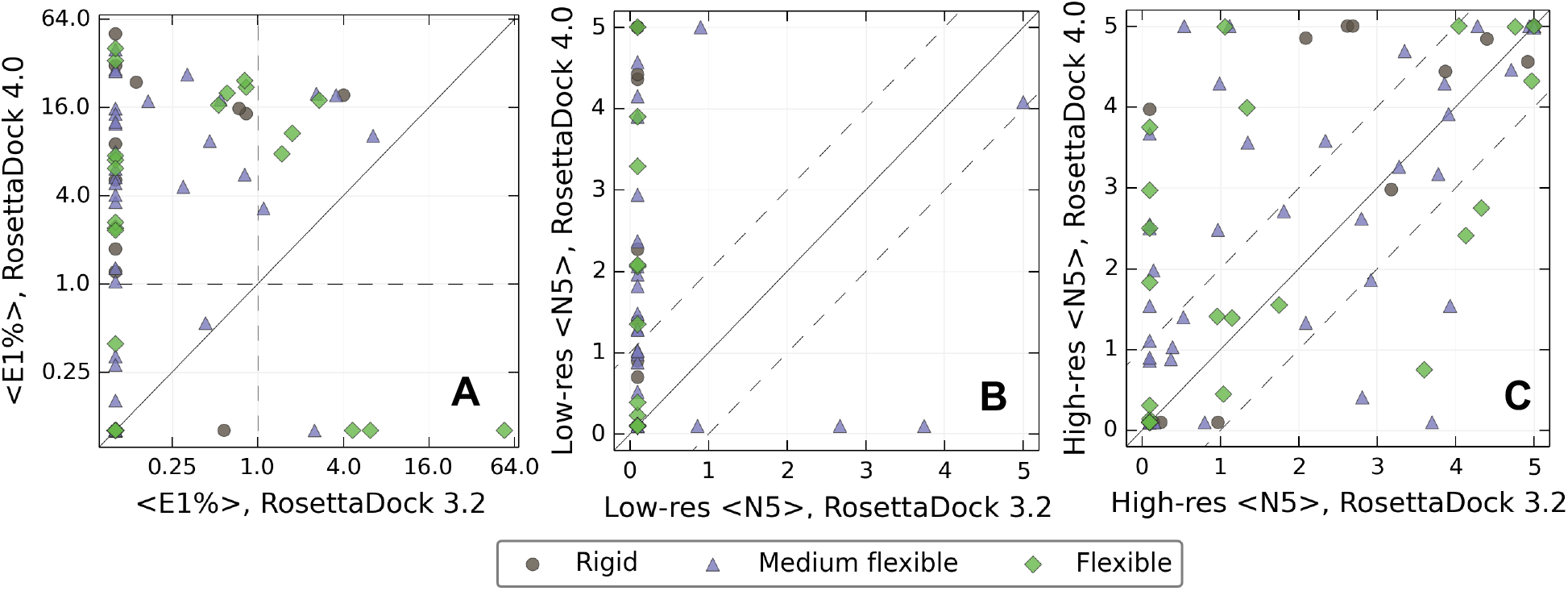
Comparison of performance metrics between RosettaDock 3.2 and RosettaDock 4.0 for individual complexes in the benchmark. Targets are represented by different symbols corresponding to their difficulty category (triangle: rigid; circle: medium; diamond: flexible). Points above the solid line represent better performance in RosettaDock 4.0, while points below the line represent better performance in RosettaDock 3.2. Comparison of (A) 〈*E_1%_*〉 enrichment values between the two protocols on a log-log axes. 〈*E_1%_*〉 shows marked improvement in the vast majority of the complexes. Dashed lines demarcate regions where the low-scoring set is enriched in near-native structures. Comparison of 〈*N5*〉 values (B) after low-resolution stage, and (C) after high-resolution stage (full protocol). Dashed lines highlight the region in which the two protocols differ significantly, *i.e*. by more than one point in their 〈*N5*〉 values. After the full protocol, 23 of the 88 complexes are modeled significantly better and 7 complexes are modeled significantly worse.

**Table 1.** Summary of performance of RosettaDock 3.2 vs. RosettaDock 4.0 across an 88-target benchmark set.

|  |  | RosettaDock 3.2 |  |  | RosettaDock 4.0 |  |  |
| --- | --- | --- | --- | --- | --- | --- | --- |
|  |  | Low-Res | High-Res |  | Low-Res | High-Res |  |
| | | $\langle N5 \rangle$ | $\langle N5 \rangle$ | $\langle E_{1\%} \rangle$ | $\langle N5 \rangle$ | $\langle N5 \rangle$ | $\langle E_{1\%} \rangle$ |
| Average Value | **Rigid Body** | 0.0 | 2.7 | 0.0 | 2.2 | 3.5 | 9.0 |
|  | **Medium** | 0.3 | 1.8 | 0.0 | 1.0 | 2.4 | 3.6 |
|  | **Difficult** | 0.0 | 1.2 | 0.0 | 0.6 | 1.6 | 0.2 |
|  | **Difficult (Doped)** |  |  |  | 0.7 | 2.2 | 2.9 |
|  | **All** | 0.1 | 1.9 | 0.0 | 1.3 | 2.5 | 2.5 |
|  | **All (Doped)** |  |  |  | 1.3 | 2.7 | 3.7 |
| Expected Successes | **Rigid Body** | 0.0 | 7.1 | 0.1 | 5.6 | 9.5 | 7.0 |
|  | **Medium** | 2.5 | 15.4 | 2.8 | 7.6 | 20.2 | 13.0 |
|  | **Difficult** | 0.0 | 7.4 | 0.3 | 3.6 | 9.8 | 5.0 |
|  | **Difficult (Doped)** |  |  |  | 4.2 | 13.7 | 4.7 |
|  | **All** | 2.5 | 29.9 | 3.2 | 16.8 | 39.6 | 25.1 |
|  | **All (Doped)** |  |  |  | 17.4 | 43.4 | 24.8 |

While the low-resolution stage is improved using binned energy approximations, additional gains are possible in the high-resolution stage where all protein atoms are explicitly represented. After the full protocol with both low- and high-resolution stages, the average 〈*N5*〉 increases from 1.9 in RosettaDock 3.2, which represents a marginal failure, to 2.5 in RosettaDock 4.0, which represents the borderline for success. The expected number of successes in the benchmark set increases from 29.9 to 39.6 complexes, a 32% improvement. About half of the additional successes are gained from moderately-flexible complexes, with another quarter coming from flexible complexes, suggesting that RosettaDock 4.0 is better at capturing flexible backbones than RosettaDock 3.2. Additionally, although rigid complexes only comprise 15% of the benchmark set, they comprise 25% of the docking improvements, suggesting that in a more balanced benchmark set containing more rigid targets, the improvement in performance in RosettaDock 4.0 might be even larger. As shown in Fig. 6C, while 23 complexes have full protocol 〈*N5*〉 values improved by 1 or more in the RosettaDock 4.0 simulations, 7 complexes have 〈*N5*〉 decreased by 1 or more. Detailed metrics for each target can be found in Supplementary Table S4 and Figs. S11-S16.

### Ensembles Doped with Near-bound Structures

We previously showed that when the RMSD gap between the closest conformation in the ensemble and the bound state exceeds 1 Å, induced fit methods are rarely able to access the binding funnel (18). We observed similar results for the Docking Benchmark 5.0 difficult targets (cases with interface RMSD_Cα_> 2.2 Å). As none of the ensemble generation methods used move the backbone quite so far, neither RosettaDock 3.2 nor RosettaDock 4.0 performed well on difficult targets. For example, the complex of SRP GTPase with FtsYh undergoes an interface conformational change of 2.67 Å RMSD (Fig. S10), and the docking run is only able to create a few acceptable predictions, but not rank them highly (Fig. 7A). Both monomer backbones undergo about 3 Å of conformational change upon binding, but the ensembles created from the unbound state do not contain any conformations closer than 2.5 Å from the bound state (Fig. 7C).

**Fig. 7.**
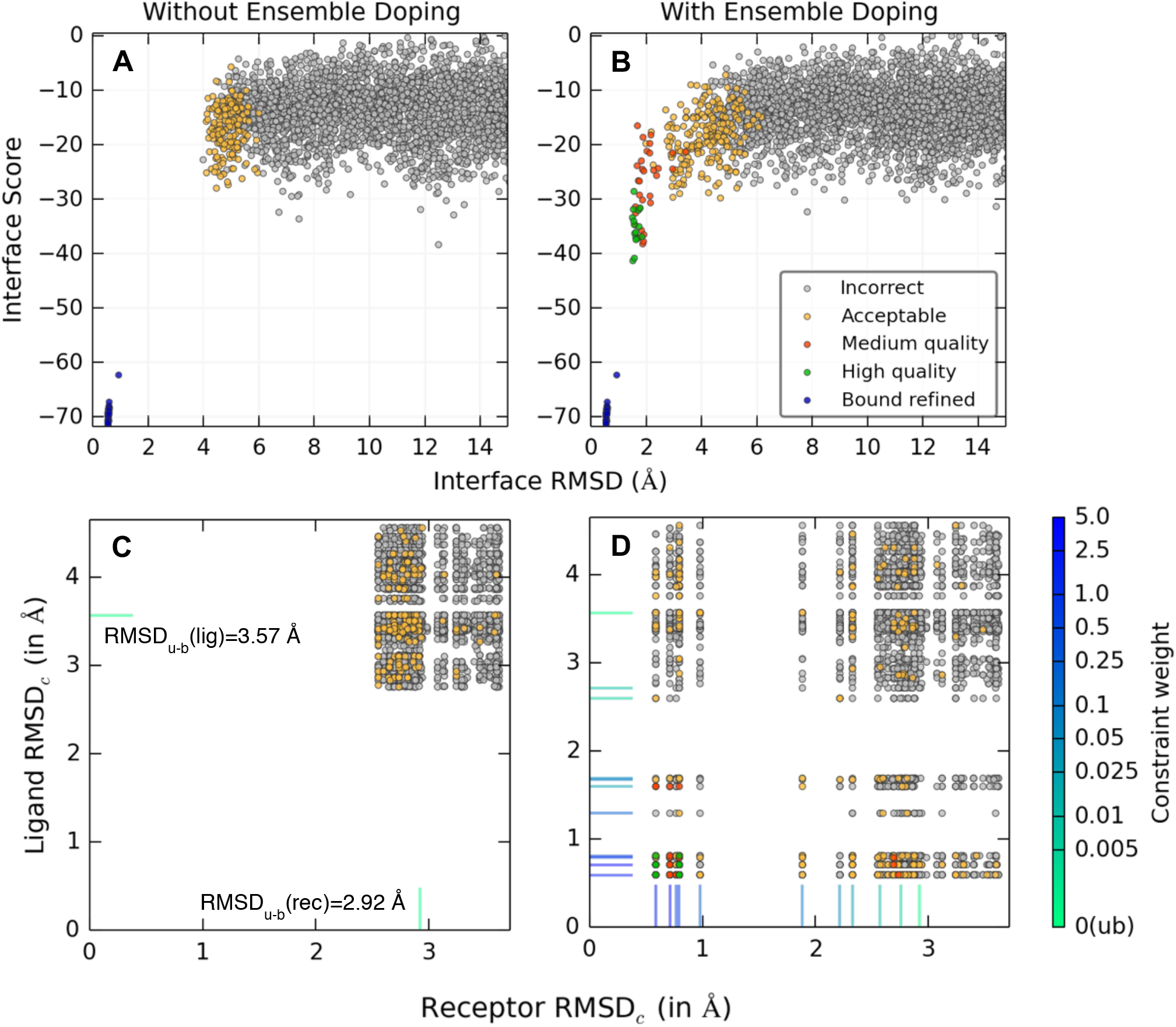
Improvement in docking performance of RosettaDock 4.0 by doping the ensemble with near-bound decoys for SRP GTPase:FtsY complex. Score versus RMSD plot of runs with (A) backbone conformations generated using NMA, Backrub and Relax protocols, and (B) ensembles doped with 10% near-bound conformations. (A) Without the ensemble doping, the simulations did not generate medium- or high-quality docked structures, and the acceptable structures did not score low enough to be discriminated from incorrect structures. (B) Ensemble doping generated deep docking funnels with high-quality structures. Colored points indicate CAPRI-quality category for each decoy, and the blue points provide a reference energy of the refined, bound crystal structure. (C and D) Plot of the contact-residue RMSD_Cα_ from the bound conformation for the ligand and the receptor conformers selected after the docking simulation for (C) ensembles without near-native doping, and (D) ensembles with 10% near-bound conformations doped. The RMSD values of the unbound conformations are marked with a green line segment, and those of the near-bound conformations are marked in colors corresponding to the biasing constraint weight. (C) The conformer generation methods are unable to generate sub-Å contact-residue RMSD_Cα_ structures starting from the unbound ligand conformation (with RMSD_Cα_ of 3.57 Å) and the unbound receptor conformation (with RMSD_Cα_ of 2.92 Å). (D) Four of the biased conformations of the ligand and five of the receptor are within 1 Å RMSD_Cα_ from the bound state. RosettaDock 4.0 is able to recognize these close conformations, find the nativelike interface and successfully dock the complex.

For cases with large backbone variation, we wondered whether RosettaDock 4.0 could select a near-bound backbone if such structures were present in a large, diverse ensemble used in the conformer selection stage. Therefore, we tested docking using ensembles doped with near-native backbone structures. To generate near-bound structures, we used Rosetta’s Relax protocol with pairwise C_α_-C_α_ distance constraints to bias the simulation towards the known bound state. Using different constraint weights, we generated 10 conformers that were progressively nearer to the bound state, with the closest 4 conformations ranging from 0.59 to 0.81 Å RMSD from the bound structure for both receptor and ligand. To complete the ensemble, we mixed these 10 structures with an unbiased set of 36 NMA structures, 27 Backrub structures, and 27 Relax structures. For the SRP GTPase:FtsY complex (PDB: 2J7P), RosettaDock 4.0 produces structures using the full range of backbone conformations (Fig. 7D). Furthermore, the lowest-scoring docked structures are near-native (Fig. 7B) and are chosen from the monomer backbones near the bound conformation (Fig 7D). Remarkably, even with just four near-bound backbones present in an ensemble of a hundred conformations with widely differing interface structures, RosettaDock 4.0 correctly recognizes these close conformations and docks them successfully. Fig. 7D shows the correlation between closer backbones and better docked structures. Similar results are seen for others including the Pol III-ε:Hot complex (PDB: 2IDO), which has a 2.79 Å interface RMSD_Cα_ between the unbound and bound states (Fig. S9). In all, the doping method was able to add nearly 4 additional expected successes among the 32 difficult targets in the benchmark set. Detailed metrics for each target can be found in Supplementary Table S4 and Figs. S17 and S18.

### Improved Efficiency for Large Ensembles

One of the principal aims was to create a protocol that scales well with increasing ensemble sizes. Fig. 8 shows run time across the benchmark set. In 77 of the 88 complexes tested, for ensembles containing 100 conformations each, RosettaDock 4.0 requires only 20–80% more time than RosettaDock 3.2 with just 1 receptor and 10 ligand conformations. Time per structure scales as 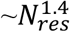 for both RosettaDock 4.0 and RosettaDock 3.2, where *N_res_* is the number of residues in the complex.

**Fig. 8.**
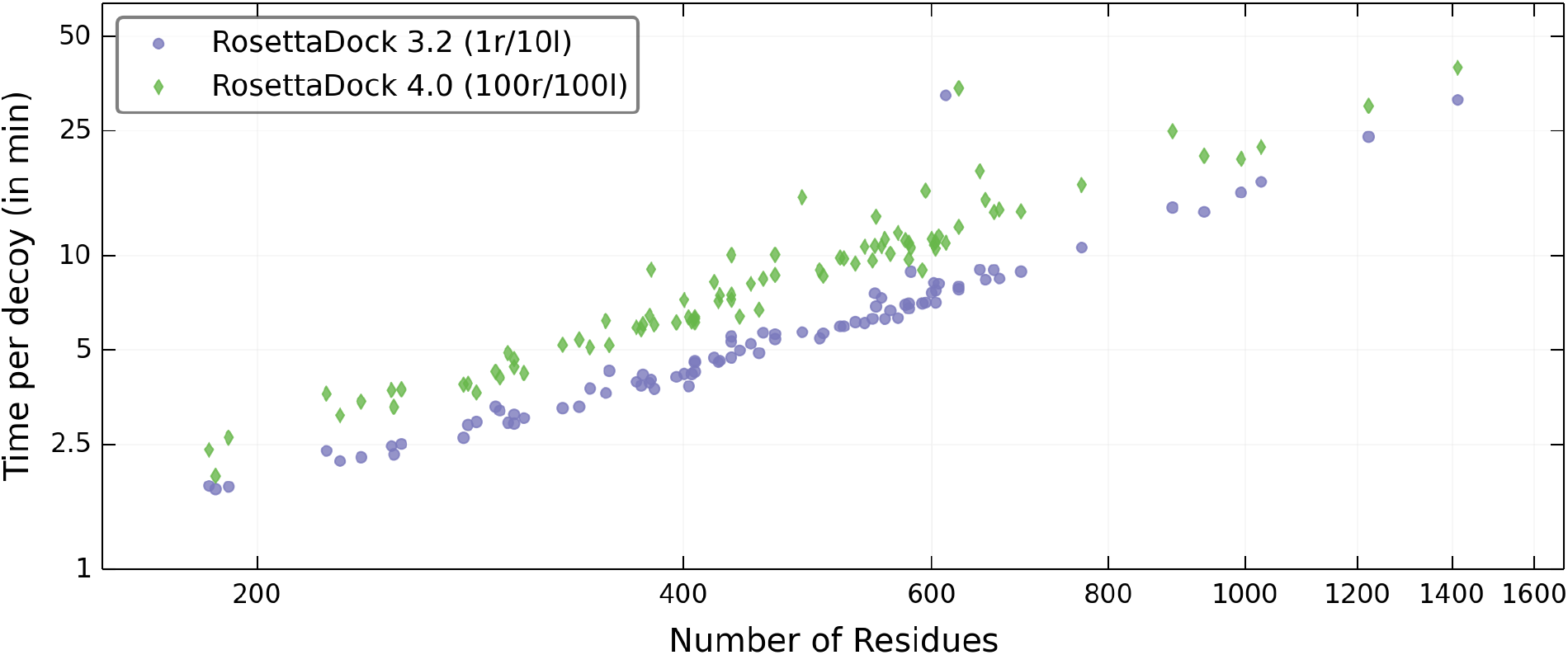
Efficiency of RosettaDock 4.0 on large ensembles. Despite sampling 100 conformations each of the receptor and the ligand as compared to 1 receptor and 10 ligand conformations in RosettaDock 3.2, the time per decoy for RosettaDock 4.0 is 20-80% more in 77 of the 88 targets tested.

## Discussion

Computational protein–protein docking can be confounded by a diverse set of problems, including backbone motion upon binding, global docking scope, and lack of structural knowledge of the docking partners. Within the Rosetta framework, all of these problems can potentially be addressed by intelligently increasing the conformational sampling space of the docking protocol. Sampling increases must be offset with efficiency gains, however, to prevent the computational costs from exploding. We developed two key advances here to create RosettaDock 4.0. First, ACS now allows us to examine a variety of backbone motions introduced by different ensemble generation protocols. The protocol scales well with an increasing number of backbones by providing adequate sampling with a runtime overhead of merely 20-80% when testing 1000-times more backbone combinations. Second, the low-resolution scoring using MDS shows a marked improvement in accuracy over centroid scoring. MDS triples the number of targets in which the top 1% of models are significantly enriched with near-bound structures, and it is seven to nine times as effective for discriminating top models, as measured by the bootstrapped 〈*N5*〉 metrics. More generally, MDS captures nearly all of the discriminatory power of the full-atom score function upon which it is based, exhibiting similar low-resolution and high-resolution *N5*, *N100*, and *N1000* metrics. Most importantly for a low-resolution score function, MDS achieves these gains in accuracy without sacrificing computational efficiency, running in roughly equivalent time to the centroid scoring method. It does require about 2 GB of additional memory to store the score table (requiring approximately 2.6 GB total compared to 0.6 GB for the baseline protocol). However, with modern computer architecture, this requirement is not prohibitive. With enhanced scoring and sampling, RosettaDock 4.0 can now select near-bound backbones in large, diverse ensembles for targets with significant changes at the interface.

RosettaDock 4.0 compares favorably to other docking protocols despite using more stringent success criteria. Table 2 summarizes recent published results from five leading docking methods: HADDOCK (45), iATTRACT (10), ClusPro (46), ZDOCK (47), and RosettaDock 3.2 (29). While the methods have different scopes and benchmarks, and report their results in different forms, we were able to assign an *N#* success metric (analogous to *N5*, *N100*, etc.) to each method. In general, current methods are good at docking easy, rigid-body targets (~50% accuracy or better), but they are all poor when the targets become more flexible (< ~30% accuracy on medium flexibility targets, <~15% on high flexibility targets). RosettaDock 4.0 maintains this level of accuracy for easy targets (77%) while showing dramatically improved accuracy for flexible targets, both among medium difficulty targets (49%) and high difficulty targets (31%). To our knowledge, this is the first report of a protein docking protocol achieving ~50% accuracy on targets with backbone flexibility between 1 Å and 2 Å RMSD. Thus, RosettaDock 4.0 marks a key step toward a paradigm shift in protein–protein docking where complexes with backbone flexibility become tractable, which has long been a goal in the community (17, 48–50).

**Table 2.** Comparison of published success rates of five leading docking methods with RosettaDock 4.0.

| Method | Description | Flexi-<br>bility? | Benchmark Set | Docking<br>Search | Success<br>Metric | Performance |  |  |  |
| --- | --- | --- | --- | --- | --- | --- | --- | --- | --- |
|  |  |  |  |  |  | All<br>Targets | Easy<br>Targets | Medium<br>Targets | Difficult<br>Targets |
| HADDOCK<br>(2017) (45) | Restraint-based<br>docking, minimization | Yes | CASP-CAPRI<br>(16) | Mixed<br>global/local | N10 = 1 | 16/25<br>(64%) | 12/12<br>(100%) | 4/13<br>(31%) |  |
| ClusPro<br>(2017) (46) | FFT docking, cluster<br>evaluation | No | CAPRI Rds.<br>13–35 | Mixed<br>global/local | N10 = 1 | 19/42<br>(45%) | 12.5†/16<br>(78%) | 6.5†/26<br>(25%) |  |
| iATTRACT<br>(2015) (10) | Rigid-body docking,<br>interface refinement | Yes | Docking<br>Benchmark 4.0<br>(3) | Global* | N200 =<br>30 | 64/166<br>(39%) | 55/119<br>(46%) | 9/28<br>(32%) | 0/19<br>(0%) |
| ZDOCK<br>(2011) (47) | FFT docking, model<br>evaluation | No | Docking<br>Benchmark 4.0 | Global | N100 =<br>1‡ | 65/176<br>(37%) | 58/121<br>(48%) | 7/30<br>(23%) | 0/25<br>(0%) |
| RosettaDock<br>3.2 (2011)<br>(29) | Monte Carlo docking,<br>model evaluation | Yes | Docking<br>Benchmark 4.0 | Local | N5 = 3 | 56/115<br>(49%) | 49/84<br>(58%) | 5/17<br>(29%) | 2/14<br>(14%) |
| RosettaDock<br>4.0 | Monte Carlo docking,<br>model evaluation | Yes | Docking<br>Benchmark 5.0<br>(4) | Local | N5 =<br>3** | 41/88<br>(47%) | 10/13<br>(77%) | 21/43<br>(49%) | 10/32<br>(31%) |

\*Nearest-native structures from rigid-body docking selected for refinement. †Half successes awarded for targets with multiple binding sites evaluated, where at least one but not all binding sites are captured. ‡2.5 Å cut-off for near-native structures. \*\*Cases where bootstrapping gives $\geq 50\%$ chance of success are included, cases where bootstrapping gives $< 50\%$ chance of success are excluded. For CAPRI sets, medium & difficult targets are combined, comprising all targets without at least one high-quality prediction by any predictor.

The limiting factor to successfully docking protein complexes with greater flexibility is now the ability to generate conformers within 0.7 Å of the bound state where MDS can start recognizing interfaces. Previously, our lab compared seven commonly used methods to generate ensembles from monomers; while ensembles from most methods had ~50% directional overlap with the experimentally observed direction, the magnitudes of these motions were insufficient to reach the bound conformations (18). Diversifying ensembles by pushing them along their top principal components may help close the gap. Moving the backbone along different combinations of principal components from different conformer generation methods could generate a larger variety of backbone conformers. Another possible solution for proteins which have been crystalized in different contexts or have structurally diverse homologs is a distance geometry-based conformer selection method, which has recently been shown to span relevant conformational space (51).

While RosettaDock 4.0 makes large strides in conformer selection, the protocol still simulates induced fit only in the all-atom mode with small, rigid-body moves and side-chain packing at the interface. This method can consistently produce high-quality docked structures only if the conformers selected are within 0.7 Å of the bound state (18). Other studies have shown significant contributions of induced fit, whether implemented via Cartesian minimization at the interface (10) or through contact-specific normal mode analysis (52). Previous attempts to introduce flexibility at the interface in RosettaDock by varying backbone torsions resulted in 3-fold increased run times for the smallest targets (53). Doing so by minimizing along Cartesian coordinates can slow the protocol down by more than 10 times (data not shown). These protocols were implemented in the high-resolution phase because the centroid score was not accurate for native discrimination. MDS might now enable induced fit methods in the low-resolution phase, adding further backbone conformer sampling. Additionally, the accuracy of MDS means that low-resolution output structures might be filtered such that only a small fraction are sent to the expensive high-resolution phase. As such, MDS will be a critical component of the future ability to the RosettaDock protocol to induce a fit at the interface.

## Methods

### PDB Curation

To create the score tables for motif dock score, we culled the Protein Data Bank (54) for all crystal structures containing two or more interacting protein chains and a resolution of 3.0 Å or better. We also removed any structures present in the Docking Benchmark 5.0 (4) to be used as a test set. We further removed all homologs of complexes in Docking Benchmark 5.0 and validated the lack of dependence on homologs (see Table S3). In the remaining set, PDB structures with more than two chains in their asymmetric unit were further divided such that one structure represented every pair of interacting protein chains in their asymmetric unit. The PDB structures were then stripped of all HETATM lines and non-canonical amino acids. Our curated set contains 154,955 protein–protein complex structures from 103,017 PDB entries.

### Motif Querying

Each structure in the protein interface set was loaded into Rosetta and scored with a full-atom score function; the resultant energies were decomposed onto the set of interacting residue pairs. The system was queried for cross-chain pairs of residues with C_β_ atoms (C_α_ for glycine) within 10 Å of each other with a pair score below a constant energy cut-off (typically 0 kcal/mol; *i.e*. residue pairs that are net-attractive). For each residue pair in the filtered residue set, we calculated the six-dimensional transform needed to superimpose one amino acid backbone onto the other (three-dimensional Cartesian translation and three-dimensional Euler angle rotation). Each pair score was stored with its corresponding 6D-transform as a one-line motif.

### Score Grid Generation

A score grid is initialized with a translational and rotational grid size. One by one, motifs are analyzed. The motif 6D-transform is binned, and the corresponding bin in the score grid is queried. If the bin is empty, the motif score is saved as the bin score. If the bin is populated, the old bin score and the motif score are compared, and the lower of the two is saved as the new bin score. If smoothing is being used, the neighboring bins are also queried, with the favorability of the score determining the radius within which the bin scores are updated (see Supplementary Method S1 for further details). Once all motifs have been analyzed, the populated bins are assigned a hash value and, to minimize its memory footprint, only the hashed bins are stored (32, 55).

### Scoring with Motif Dock Score

RosettaDock 4.0 uses the same algorithmic framework as RosettaDock 3.2 described previously (34), with modernizations described in thereafter (13, 29, 30). The standard low-resolution score function (`interchain_cen`) is replaced with a motif-based score function, called `motif_dock_score`. The score function consists of a new scoring term, `motif_dock`, and a clash penalty (`interchain_vdw`). The `motif_dock` term is a residue pair energy that acts only on cross-chain residue pairs with C_α_ atoms within 10 Å of each other. The residue pairs are scored by calculating their 6D-transform, converting this to the hash value of the corresponding 6D bin, querying the hash table, and reporting the bin score. If the bin is empty (i.e. there are no matches for the hash), the pair score will either be zero if no penalty is used, or 0.5 kcal/mol, if a penalty is used.

### Benchmark Set Generation

We built three benchmark sets using subsets of the Docking Benchmark 5.0. The first, a set of eleven targets for rescoring, was randomly selected from the rigid-body subset of Docking Benchmark 5.0 to provide ample near-bound structures to optimize motif scoring’s near-native discrimination ability. To generate the rescoring sets for each target, we ran the standard RosettaDock protocol (29) on the unbound complex structures, including translation and rotation perturbations (mean = 3 Å translation, 8° rotation) to the ligand (the smaller protein partner) to disrupt existing interfaces. The second set, a small representative docking benchmark, was generated by selecting four rigid targets (1EFN, 1GLA, 2A1A, 2FJU), three medium-flexibility targets (1LFD, 2CFH, 3AAA), and two flexible targets of different categories (2OT3, 3F1P) from the Docking Benchmark 5.0. The third set, a larger representative docking benchmark, contained all targets in the second benchmark, as well as all rigid-body targets tested previously (13), which still remained in Docking Benchmark 5.0 (13 in total), all remaining medium difficulty targets from Docking Benchmark 5.0, and 32 additional difficult targets chosen randomly from the Docking Benchmark 5.0 set. (We could not generate ensembles for the receptors of 1N2C, 3R9A, 1DE4 and 4GAM in reasonable time that would otherwise have been included in our benchmark set.)

### Generation of Backbone Ensembles

To generate diversity in backbone conformations for the RosettaDock 4.0 runs, we used three conformer generation methods: perturbation of the backbones along the normal modes by 1 Å (44) (using RosettaScripts (56)), refinement using the Relax protocol in Rosetta (42), and backbone flexing using the Rosetta Backrub protocol (43). (Detailed protocols and complete command lines are provided in Supplementary Method S2.) Each was shown to generate different modes of backbone motions, overlapping by 30–50% with the actual directions of motion between the unbound and bound states (18). Since the normal mode analysis generated the largest deviations, we used 40 normal mode conformers, 30 Relax conformers and 30 Backrub conformers to comprise the ensemble of 100 conformers.

### Doping Ensembles with Near-bound Structures

For difficult targets, the three ensemble generation methods often fail to generate near-bound conformations. Thus, we also evaluated the ability of RosettaDock 4.0 to discriminate bound-like structures with the help of conformations biased to resemble the bound state. Starting with the unbound conformation, we relaxed the structure while employing C_α_ distance constraints for all pairs of residues (except for adjacent residues). These C_α_ constraints are harmonic potentials with the mean distance set to the corresponding C_α_–C_α_ distances in the bound conformation and a spring constant of 1 kcal/mol/Å^2^. To create a library of intermediate structures, we set the weight of the constraints to 0.005, 0.01, 0.025, 0.05, 0.1, 0.2, 0.5, 1.0, 2.5, and 5.0. The ensembles were then doped with the 10 intermediate structures after proportionally removing 10 structures from the existing ensemble.

### Docking Simulations

Docking simulations were performed using two versions of RosettaDock, *viz*. 3.2 (13) and 4.0 (developed in this article). Briefly, RosettaDock is a Monte Carlo-plus-minimization-based docking algorithm, which works in two stages. The first is a coarse-grained, low-resolution stage where side chains are represented as pseudo-atoms. The smoother energy landscape of this stage allows large rigid body motions and backbone conformer selection to occur simultaneously. It is in this stage that both the sampling and scoring enhancements have been implemented in version 4.0. The second all-atom, high-resolution stage is common to both versions. In this stage, finer rigid body motions are accompanied by interface side-chain packing and energy minimization in torsion angle space (ϕ, ψ and χ_i_). (Detailed protocols and complete command lines are provided in Supplementary Method S3.)

### Benchmark Evaluation and Success Metrics

Docking runs with *N#* values above a given threshold are categorized as “successful”. For *N5*, we define 3 near-native decoys as a success when evaluating docking protocols. We also use *N50*, *N100*, *N500*, and *N1000* (success thresholds of 15, 30, 75, and 150, respectively) to measure the sampling rates of near-native models in our top 1% and top 10% of models, respectively. We use *E_1%_* and *E_10%_* to measure the ability of our scoring methods to enrich a model set. We calculated the expected value of *N#* and *E_N%_* metrics by sampling with replacement from the available model set a number of models equal to the size of the set. This was repeated 1000 times and averages calculated from these calculations are denoted by 〈·〉.

To be counted as near-native, our high resolution models must meet the standard criteria for a CAPRI acceptable, medium-quality or high-quality model (i.e. have f_nat_ ≥ 0.1 and, either ligand RMSD ≤ 10.0 Å or interface RMSD ≤ 4.0 Å) (57). Here, f_nat_ is the ratio of the number of native residue-residue contacts in the predicted complex to the number of contacts in the experimental structure of the bound complex, ligand RMSD is the backbone RMSD of the ligand molecule in the predicted complex versus the experimental complex upon receptor superposition, and interface RMSD is the interface heavy-atom RMSD after superposition of the backbone atoms of the interface residues. We use a more lenient measure for our low-resolution decoys (centroid RMSD ≤ 6.0 Å) to account for the limitations of measuring RMSD in the centroid phase (incompletely resolved side chains, lever-arm effects away from the interface etc.).

## Acknowledgements

This work has been supported by the National Institutes of Health, USA (grant R01-GM078221). Computations in this study have been performed in part on the Maryland Advanced Research Computing Center (MARCC) cluster.

## Availability

As a part of the Rosetta software suite, RosettaDock 4.0 is available to all noncommercial users for free and to commercial users for a fee.

## Author contributions

N.A.M., S.S.R.B., and J.J.G. designed research; N.A.M., and S.S.R.B. performed research; W.S. contributed underlying architecture for MDS; N.A.M., S.S.R.B., and J.J.G. analyzed data and wrote the paper.

## Conflict of Interest

J.J.G. is an unpaid board member of the Rosetta Commons. Under institutional participation agreements between the University of Washington, acting on behalf of the Rosetta Commons, Johns Hopkins University may be entitled to a portion of revenue received on licensing Rosetta software, which includes the methods described in this paper.

